# Rottlerin ameliorates DSS-induced colitis by improving intestinal barrier function via activation of the Epac-2/Rap-1 signaling pathway

**DOI:** 10.1101/2020.03.16.994582

**Authors:** Xue Song, Lugen Zuo, Luyao Wang, Zihan Zhu, Jing Tao, Yifan Jiang, Xiaopei Wu, Zhikun Wang, Jing Nian, Ping Xiang, Xiaofeng Zhang, Hao Zhao, Liang Yu, Jing Li, Jianguo Hu

## Abstract

**OBJECTIVES:** Rottlerin, a pan PDE inhibitor, has a variety of pharmacological activities, including enhancing barrier function and mediating anti-inflammatory activity by changing the distribution of occludin and ZO-1. Nevertheless, the function of rottlerin on Crohn disease (CD) keep unknown. Our aim of the study is to investigate the role of rottlerin on CD-like colitis and its mechanism.

**METHODS:** Wild-type mice which were 8-10 weeks old were randomly divided into three treatment groups: (i) the normal feeding, no administration (control) group, (ii) the group administered 3% dextran sodium sulfate (DSS) alone, and (iii) the group administered rottlerin (100 mg/kg) and 3% DSS. In this study, the effect of rottlerin on the function and structure of the intestinal barrier was investigated, and the possible mechanism was discussed. We performed signaling pathway analysis and flow cytometry to identify the detailed mechanisms by which rottlerin (10 μg/mL) treatment inhibits cell growth arrest and the attenuation of TJ proteins in LPS-treated FHs 74 int cells.

**RESULTS:** Rottlerin treatment significantly ameliorated colitis induced by DSS in WT mice, which was manifested by a decrease in inflammation score, the attenuation of inflammatory factors and the inhibition of destruction on intestinal barrier structure. Rottlerin enhanced the levels of occludin and ZO-1, and improved the function of intestinal barrier, which may have been why rottlerin ameliorated colitis in WT mice. The anti-inflammatory effect of rottlerin may be partly due to the activation of Epac-2/Rap-1 signaling.

**CONCLUSIONS:** Rottlerin may treat CD in humans via enhancing TJ proteins expression and improving the function of intestinal barrier.

## 1. Introduction

CD is considered to be a kind of chronic intestinal inflammation that is common in gastrointestinal diseases ^[1]^. CD is characterized by recurrent diarrhea, bloody stool and malnutrition and seriously harms the physical and mental health and quality of life of patients ^[2]^. Long-term inflammatory stimulation also raises the risk of colorectal cancer, causing grievous social and medical economic burdens. In recent years, with the continuous improvement of the economy, the incidence of CD has increased significantly ^[3]^. The etiology and pathogenesis of CD are not completely clear, but intestinal mucosal immune homeostasis disorder, intestinal barrier dysfunction, bacterial infection and genetics are generally recognized as factors ^[4]^.

Changes in intestinal barrier function have a significant impact in the progression and recurrence of CD ^[5]^. Intestinal barrier dysfunction causes intestinal bacteria to enter the body through the mucosa, causing local or systemic inflammatory responses ^[6]^. Since the completeness of epithelial barrier is important to resist the entry of antigens and maintain the stability of the internal environment, the primary therapeutic goal of CD is to induce and maintain healing of the intestinal mucosa ^[7]^. Effectively regulating intestinal mucosal barrier function is of great significance to improve the prognosis of CD and the quality of life of patients.

The epithelial barrier is an integral part of mucosal barrier, which protects against the transmission of macromolecular substances ^[8]^. The function of intestinal barrier is realized through various compositions, containing the epidermis made up of epithelial cells that are plugged through tight junctions (TJs) at the cell boundary seam, mucogel layers, and antimicrobial peptides ^[9]^. Several bacterial pathogens can alter TJs by altering the distribution of occludin and ZO-1. The abnormal expression of TJ proteins in the epithelial cell monolayer may disrupt the structure and function of TJs, thereby destroying the completeness of barrier ^[10]^. Inflammation is considered to be the basic mechanism of the pathophysiology of intestinal damage ^[11]^. Relative increases in the levels of inflammatory factors (TNF-α, IL-17A and IL-1β) may attenuate the expression of TJ proteins, leading to intestinal barrier dysfunction and promoting colonic mucosal inflammation ^[12]^.

Rottlerin is a polyphenolic substance that is abundant in a tropical plant from India; recent studies have shown that it have an anti inflammatory action, meanwhile is a promising medicine for the treatment of arthritis through the attenuation of phosphodiesterases (PDEs) ^[13]^. Among PDEs, PDE-4 is thought to inhibit a variety of inflammatory cytokines ^[14]^. Recent studies have shown that rottlerin has a strong ability to inhibit the decrease in transepithelial electrical resistance and improve barrier function by enhancing the distribution and levels of occludin and ZO-1 ^[15]^. Moreover, the inhibition of PDE-4 not only inhibits MAPK, NF-κB, PI3K/mTOR, and JAK/STAT-3 activation but also activates the Epac-2/Rap-1 pathway ^[16]^. Rap-1 is involved in OXPAPC-induced barrier enhancement and cell-ligation rearrangement ^[17]^. Rap-1 is activated by extracellular signals through various regulatory proteins, such as Epac-2, which may have a significant impact in various processes, including the managing of growth and differentiation, cell adhesion, and the enhancement of barrier function ^[18]^. All of these data indicate that rottlerin is an ideal potential therapeutic drug for CD, and its safety makes this medicine particularly potential.

The DSS-induced chronic colitis mouse model is considered to have the closest phenotype to CD ^[19]^. This study investigated the therapeutic effect and mechanism of rottlerin on DSS-induced CD colitis in Wild-type (WT) mice. The expression of CD, PDE-4, Epac-2 and Rap-1 was abnormal in the colon, and rottlerin not only inhibited the expression of the PDE-4 isoform but also activated the Epac-2/Rap-1 pathway, thereby increasing the levels of ZO-1 and occludin. These results provide evidence for rottlerin as a drug for the treatment and inhibition of CD.

## 2. Materials and methods

### 2.1 The preparation of patient specimen

The detection was permitted by the institutional review board, and got informed consent from patients. Intestinal samples were gathered from CD patients (n = 12), and intact intestine was gathered from the patients with colorectal carcinoma (control, n = 16). The CD patients participating in the detection (4 males and 8 females; age: 36.5 ± 4.1 years old; BMI: 16.8 ± 0.8 kg/m^2^) were classified as A2; L3; B2 by Montreal classification. Patients with colorectal carcinoma (10 males, 6 females; age: 61.2 ± 6.1 years; BMI: 16.3 ± 0.6 kg/m^2^) were classified as control sample.

### 2.2 Animals

All operation about experimental animal were permitted by the local Animal Ethics Committee. WT mice were purchased from the America Jackson Laboratory and raised in local animal experimental center (Bengbu, China). All practices about animals were executed entirely accordance with the National Institutes of Health (NIH) guidelines for animal care and use.

### 2.3 Drug administration

WT mice (male, ten weeks old) were randomly allocated to three groups (the normal group, the vehicle group, which received only DSS (dextran sulfate sodium), and the treatment group, which received DSS + rottlerin) with 10 mice per group. DSS was dissolved in drinking water, the colitis was induced by DSS (MP Biomedicals, Cat #: 160110) for 2 circulations. For the first circulation, DSS was administered for 4 days, and then normal drinking water was given for another 10 days; based on previous reports ^[19]^, the oral administration of rottlerin (100 mg/kg, Sigma Aldrich, Cat #: R5648) began after the administration of DSS in the first circulation ^[20]^. Rottlerin was dissolved as a suspension in 0.9% NaCl (Sigma Aldrich, Cat #: 793566) and orally administered once daily until the end of the second circulation (the steps of the second circulation were the same as those of the first). After 4 weeks, the mice were humanely sacrificed and the entire colon of each mouse was collected.

### 2.4 Cell culture and treatment

The human intestinal epithelial cell line FHs 74 int (ATCC CCL-241) was purchased from the Beijing Institute of Oncology. The cells were cultured in Hybri Care medium (ATCC 46X) reconstituted following the instructions of the ATCC and contained 10% FBS (Sigma, Cat #: 12007C) and 45 ng/mL epidermal growth factor (Sigma, Cat #: E5160). For execution experiments, cells were inoculated in 6-well cell culture cluste (4 × 10^5^ cells/well). Cells were cultured for 24 hours with (1) control (normal DMEM); (2) LPS treatment (10 µg/mL LPS) ^[21]^; (3) LPS with rottlerin (dissolved in DMSO, Sigma Aldrich, Cat #: D4540) treatment (10 µg/mL LPS and 1 μM rottlerin) ^[15]^; or (4) LPS with rottlerin and HJC0197 (Epac inhibitor, dissolved in DMSO, APExBIO, Cat #: 1383539-73-8) (10 µg/mL LPS, 1 μM rottlerin and 10 μM HJC0197) ^[22]^.

### 2.5 Colitis symptom assessment

All the WT mice involving this study were scored once two days with a numerical disease activity index (DAI) as reported by Spencer et al ^[23]^.

### 2.6 Histological analysis

The colon tissues of mice were paraffin-embedded for conventional dyeing with hematoxylin and eosin (H&E) and follow up pathology analysis. The sections were graded according to intestinal inflammation as reported by Schultz et al ^[24]^.

### 2.7 Intestinal permeability *in vivo*

At the end of therapeutic schedule, mice were stopped serving food for 4 hours and then gavaged with fluorescein isothiocyanate (FITC)-dextran (Sigma, Cat #: F-7250) as previously reported ^[6]^.

### 2.8 Bacterial translocation

Bacteria were derived from the tissue of mesenteric lymphnodes (MLNs) and liver for isolating via aseptic techniques as previously reported ^[6]^.

### 2.9 TUNEL staining

As previously reported ^[25]^, apoptotic cell of colon tissues was analysed and counted by TUNEL staining.

### 2.10 Enzyme-linked immunosorbent assay (ELISA)

The protein existed in the lysates of proximal colon tissues were extracted from the frozen samples, and the protein levels of IL-1β, TNF-α and IL-17A were detected using the ELISA kits (R&D Systems, Emeryville, Cat #: MLB00C, MTA00B, and M17AF0).

### 2.11 Cell proliferation assays and flow cytometry (FCM)

After 24-hours cultivation, FCM was performed to estimate the proportion of cellular apoptosis with Annexin V-FITC/PI apoptosis detection kit (Dalian Meilunbio Company, Dalian, China). The cells in the quadrants on the right of all the graphs were defined as apoptotic cells. Cells were fixed with 70% acetic acid ethanol (1 h at 4°C) for analysis of the cell cycle, resuspended in 100 µl of propidium iodide solution (PI, Sigma Aldrich, Cat #: P4170) and cultured in the dark for 30 min. A FACSCalibur flow cytometer (BD, USA) was used to define the proportion of cells in every stage of generation cycle.

### 2.12 Western blot analysis

Western blotting was executed to determine changes in protein levels of bowel as previously described ^[26]^. Primary antibodies as antibodies against Bcl-2, Bax, cleaved caspase-3, p-PI3K, p-Akt, p-mTOR, ZO-1, occludin, PDE-4, Epac-2, Rap-1 and β-actin (Abcam; Cat #: ab182858, ab32503, ab214430, ab182651, ab179463, ab109268, ab216327, ab151549, ab251568, ab14628, ab21238, ab175329 and ab81283, respectively) were used, the dilution ratio is 1:800.

### 2.13 Immunofluorescence and immunohistochemical analysis

Immunofluorescence was executed on colonic tissue using antibodies against ZO-1 and occludin (1:100; Abcam) to determine the changes in TJ protein levels of bowel. Immunohistochemistry was executed on human colonic tissue using antibodies against Epac-2 and Rap-1 (1:100; Abcam).

### 2.14 qRT-PCR

qRT-PCR analysis of intestinal proinflammatory factors, including TNF-α, IL-1β, and IL-17A, was performed as previously described to evaluate changes in mRNA levels ^[27]^.

**Table 1.**
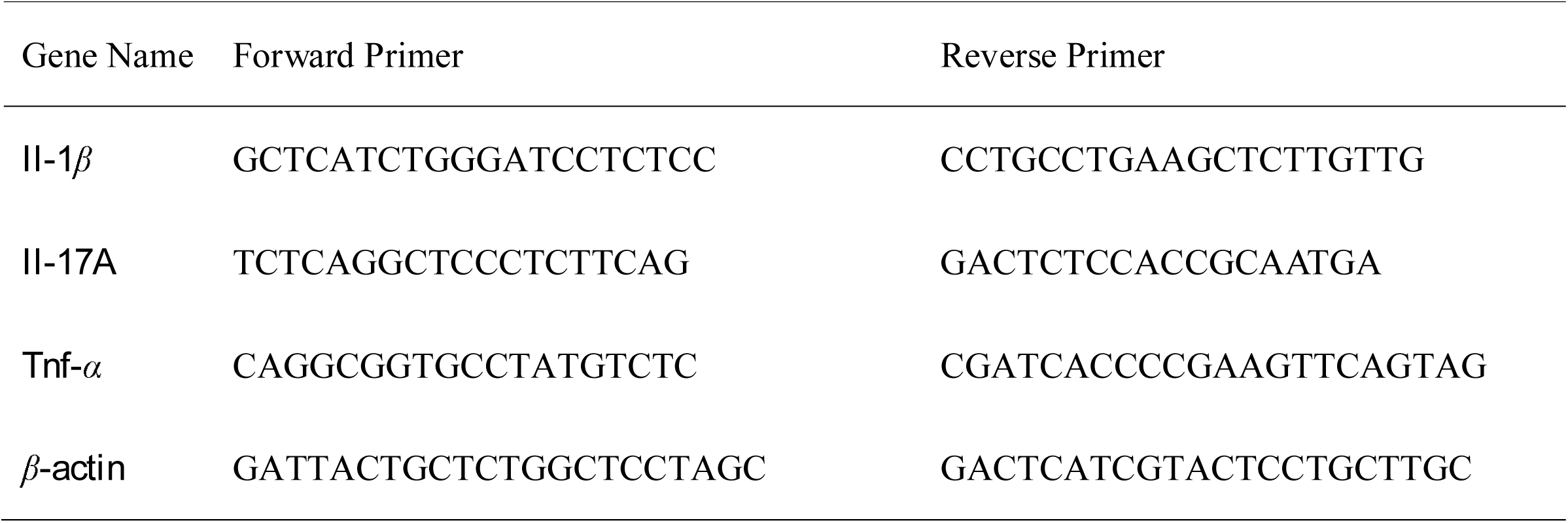
Primer Sequences (5’ to 3’)

### 2.15 Statistical analysis

The experimental data were analyzed by SPSS 17.0 software (SPSS Inc., Chicago, IL). Data from two groups were evaluated by the independent-samples t-test and categorical data were evaluated by Chi-squared tests. When *P* value was less than 0.05, it was statistically significant.

## 3. Results

### 3.1 Effect of rottlerin on DSS induced colitis in WT mice

The severity of colitis was significantly reduced over the eighteen days after administration with rottlerin. Meanwhile, the results of DAI scores exhibited the values of the rottlerin group reached levels lower than those of the DSS group (**Figure 1A**). In addition, the inflammatory scores of intestinal tissues in rottlerin group were below the scores of the DSS group **(Figure 1B-C**). Compared with the DSS group, the levels of inflammatory markers such as TNF-α, IL-1β and IL-17A in colon tissues of rottlerin group were significantly reduced (**Figure 1D**). The results of qRT-PCR further showed that the levels of TNF-α, IL-1β and IL-17A mRNA in the colon tissues of rottlerin group were decreased (**Figure 1E**). The TUNEL assay exhibited that the amount of positive cells in each crypt of the rottlerin group was below that of the DSS group, but similar to the amount of positive cells in the WT group (**Figure 1F-G**). Bcl-2 as an anti apoptotic factor, its expression level was raised in the rottlerin team compared to the DSS-treated team; however, the level of Bcl-2 in rottlerin team was still below that in WT mice (**Figure 1H-I**). In contrast, the protein levels of caspase-3 and Bax in the rottlerin the group were below those in the DSS-treated group; however, the level of them in rottlerin team were still higher than that in WT mice (**Figure 1H-I**). Some of these findings proved that colitis and the apoptosis of epithelial cell were decreased by the rottlerin treatment, rottlerin exerted a protection function on DSS-induced mice.

**Figure 1.**
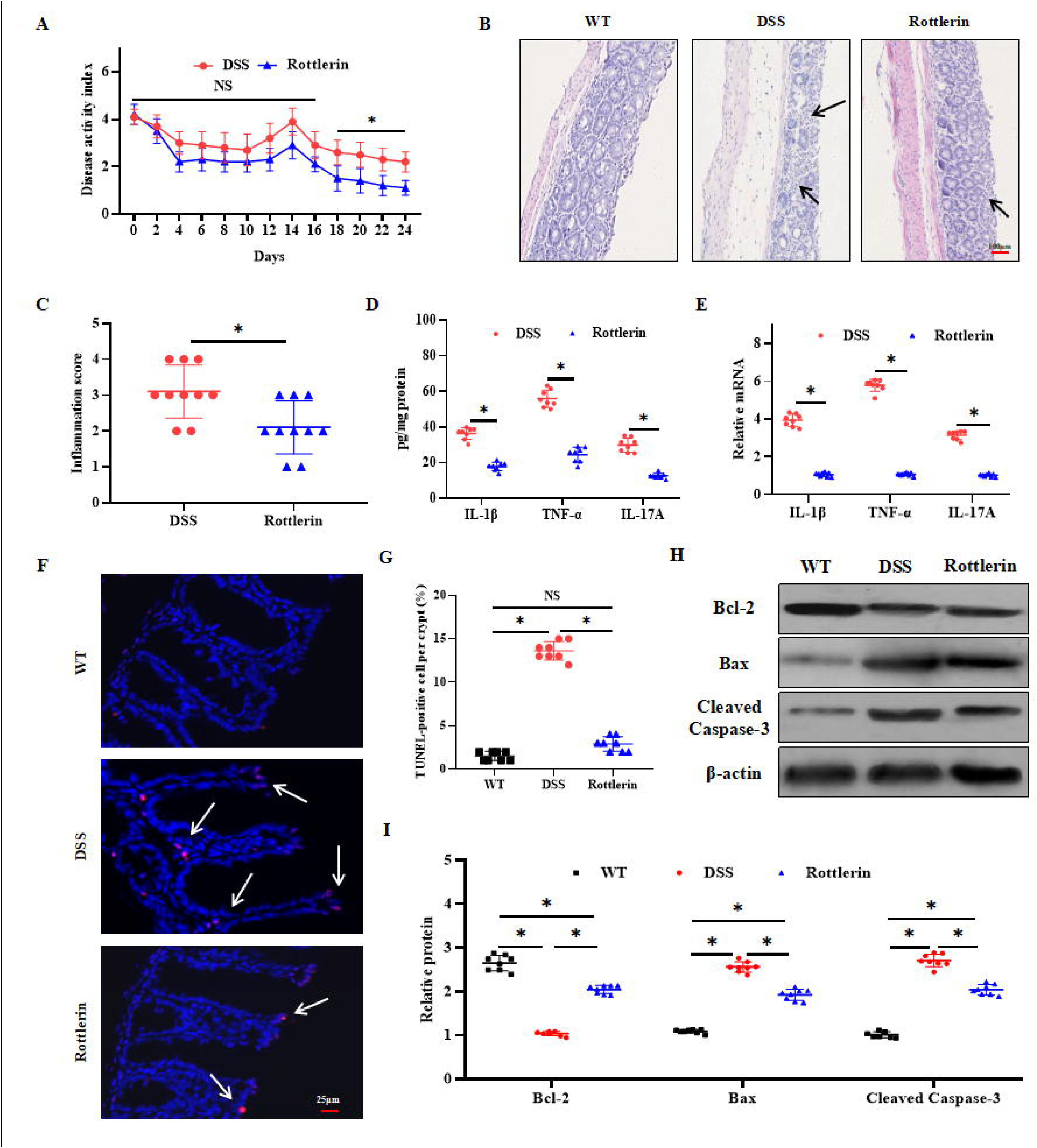
Effect of rottlerin on DSS induced colitis in WT mice. (A) The results of DAI scores of the rottlerin-treated mice were below those of the DSS-treated mice over eighteen days after treatment. (B) H&E staining revealed that rottlerin treatment alleviated the histological appearance of induced colitis. The inflamed area was indicated by arrow. (C) Inflammatory scores of intestinal tissues showed that the rottlerin treatment obviously weakened inflammation in colonic tissue gathered from the rottlerin-treated mice in contrast to the DSS-treated mice. (D) The levels of inflammatory markers such as TNF-α, IL-1β and IL-17A were obviously weakened in colonic tissue gathered from the rottlerin group in contrast to the DSS-treated group. (E) This trend was confirmed again by the results of qRT-PCR. (F-G) The TUNEL assay exhibited that the amount of positive cells in rottlerin group was below the DSS group. The positive cells were indicated by arrow. (H-I) The expression level of Bcl-2 was raised in the rottlerin team in contrast to the DSS-treated team but below that in WT mice. However, the expression levels of Bax and cleaved caspase-3 appeared an opposite trend in the rottlerin group. The detection were executed 3 times separately (8 mice of every group), a representative result is given. The datas are appeared as the means ± SD (\**P* < 0.05).

### 3.2 Rottlerin treatment improved the distribution of TJ proteins and reduced the permeability of the intestine induced by DSS

In order to investigatet whether the distributionand and levels of TJ proteins have changed after rotolin treatment, immunofluorescence and Western blotting were performed to detect the levels of occludin and ZO-1 protein in the colon of mice. The levels of occludin and ZO-1 protein in the colon of rottlerin group were dramatically upregulated in contrast to the DSS-treated group. However, the levels of occludin and ZO-1 protein in the rottlerin group were still below those in the WT mice (**Figure 2A-B**). This trend was confirmed again by the results of western blotting (**Figure 2C-D**). Intestinal permeability in mice was detected, the datas suggested that serum glucan binding levels in the rottlerin team were below those in the only DSS-treated mice, but similar to the WT mice (**Figure 2E**). Bacteria extracted from mln and liver tissue are cultured by aseptic technology. In the rottlerin group, the probability of bacterial transfer to liver and MLN was below the DSS-treated group, but similar to the WT mice (**Figure 2F-G**). All these findings partly demonstrate that intestinal mucosal barrier damage were inhibited by the rottlerin treatment in DSS-induced mice.

**Figure 2.**
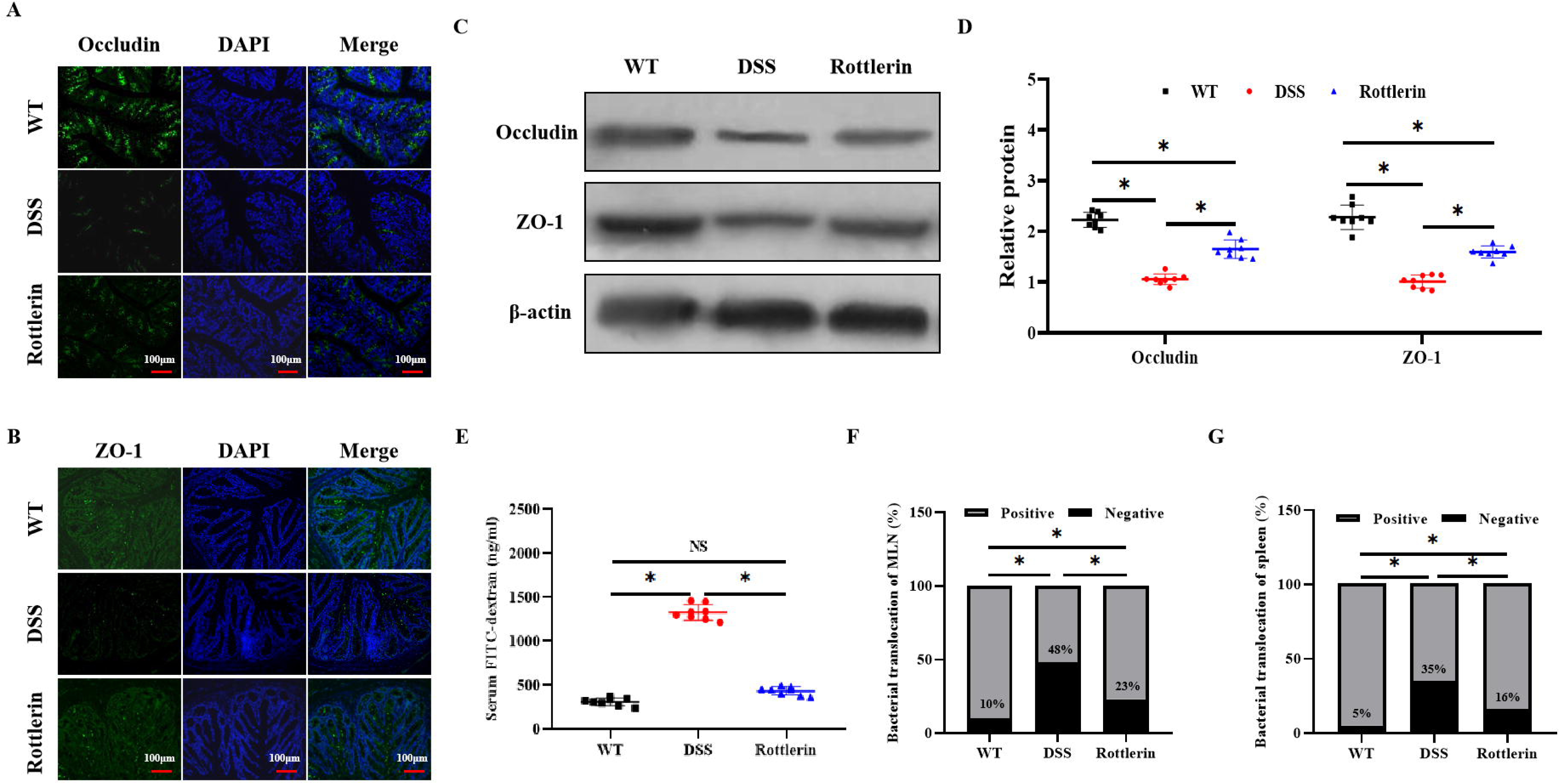
Rottlerin treatment improved the distribution of TJ proteins and reduced the permeability of the intestine. (A-B) IF revealed that the distributions of occludin and ZO-1 (green) in the colon of rottlerin group were dramatically upregulated in contrast to the DSS-treated group. The nuclei was stained by DAPI (blue). (C-D) This trend was confirmed again by the results of western blotting. (E) Intestinal permeability in mice was detected, the datas suggested that serum glucan binding levels in the rottlerin team were below those in the only DSS-treated mice, but similar to the WT mice. (F-G) In the rottlerin group, the probability of bacterial transfer to liver and MLN was below the DSS-treated group, but similar to the WT mice. The detection were executed 3 times separately (8 mice of every group), a representative result is given. The datas are appeared as the means ± SD (\**P* < 0.05).

### 3.3 Rottlerin treatment induced the activation of Epac-2/Rap-1 and the attenuation of the PI3K/Akt pathway in DSS-treated mice

Western blotting for PDE-4, Epac-2, Rap-1, p-PI3K, p-mTOR and p-Akt was executed on colonic tissues gathered from mice. The datas showed that p-PI3K, p-mTOR and p-Akt were barely expressed in colonic tissues gathered from WT mice. However, the expressions of p-PI3K, p-mTOR and p-Akt were upregulated in the DSS-induced mice but were obviously weakened after administration with rottlerin (**Figure 3A-B**). Interestingly, we found that Epac-2 and Rap-1 were highly expressed in the colonic tissuesgathered from WT mice. However, both were undetectable in the DSS-treated group not administered rottlerin but were obviously activated in the rottlerin group; PDE-4 showed the opposite trend (**Figure 3C-D**). Some of these findings proved that rottlerin promoted the reduction of inflammation and that the activation of Epac-2/Rap-1 had some relationship with this result.

**Figure 3.**
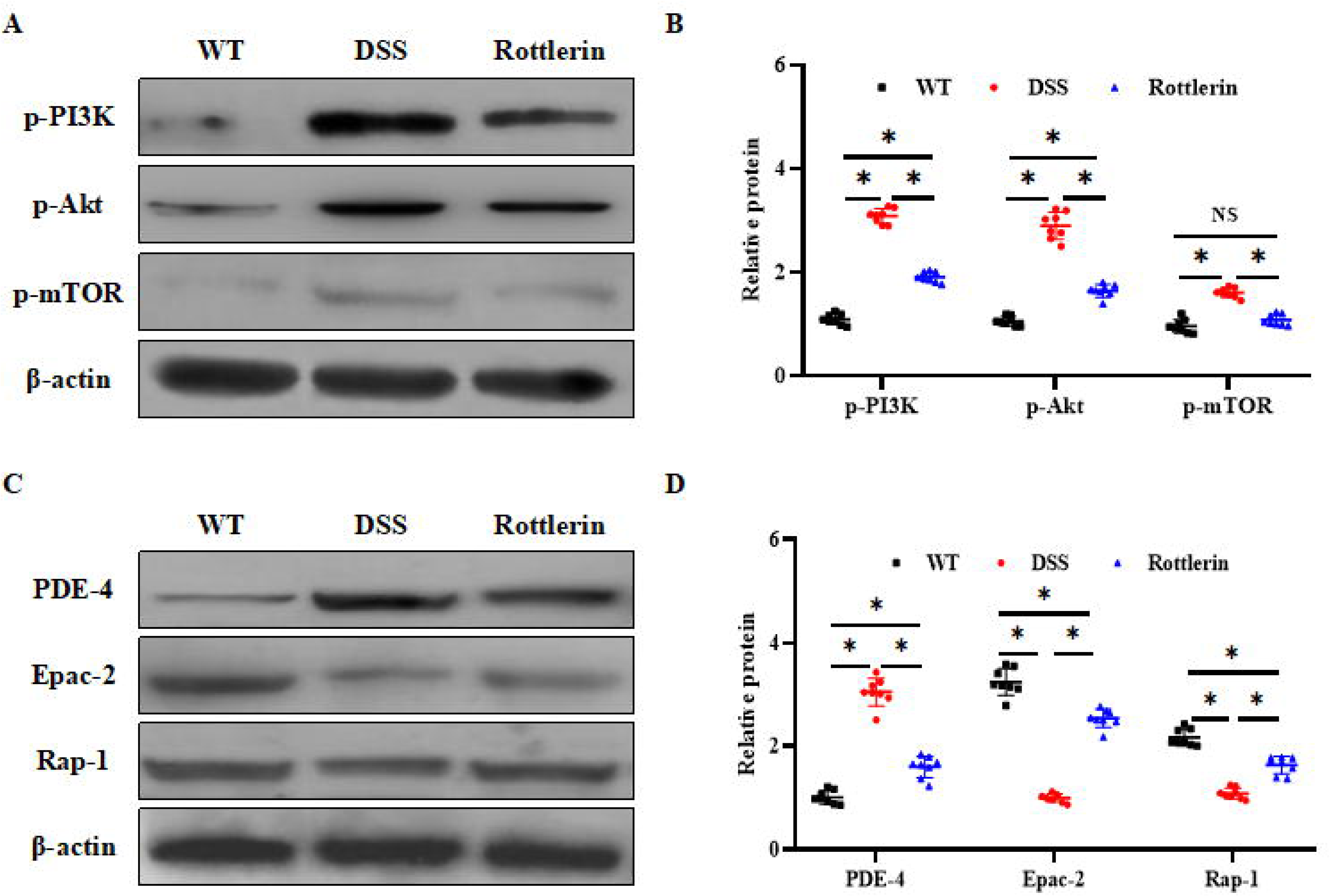
Rottlerin treatment attenuated PI3K/Akt signaling and activated the Epac-2/Rap-1 pathway. (A-B) Western blot analysis showed that the expressions of p-PI3K, p-mTOR and p-Akt were upregulated in the DSS-induced mice but were obviously weakened after administration with rottlerin. (C-D) Western blot analysis showed that Epac-2 and Rap-1 were undetectable in the DSS-treated group not administered rottlerin but were obviously activated in the rottlerin group; PDE-4 showed the opposite trend. The detection were executed 3 times separately (6 mice of every group), a representative result is given. The datas are appeared as the means ± SD (\**P* < 0.05).

### 3.4 The expressions of Epac-2 and Rap-1 decreased in the colonic tissue of CD patients

With the aggravation of inflammatory reactions, the expression of Epac-2 and Rap-1 decreased. We found the expressions of Epac-2 and Rap-1 were significantly attenuated in the colonic tissue of CD patients in contrast to the control patients (**Figure 4A-B**). We as well analyzed the relationship between the parameter of CD patients and Epac-2 and Rap-1 expression, but we didn’t find a clear connection. Western blotting for Epac-2 and Rap-1 was performed on intestinal tissue samples extracted from patients. The results showed that the levels of Epac-2 and Rap-1 were as well lower in inflammation focus of intestinal tissues from CD patients in contrast to the uninflamed intestines (**Figure 4C-D**). The attenuated Epac-2 and Rap-1 expression in the colonic tissue of CD patients suggests that Epac-2 and Rap-1 may be associated with the progression of CD.

**Figure 4.**
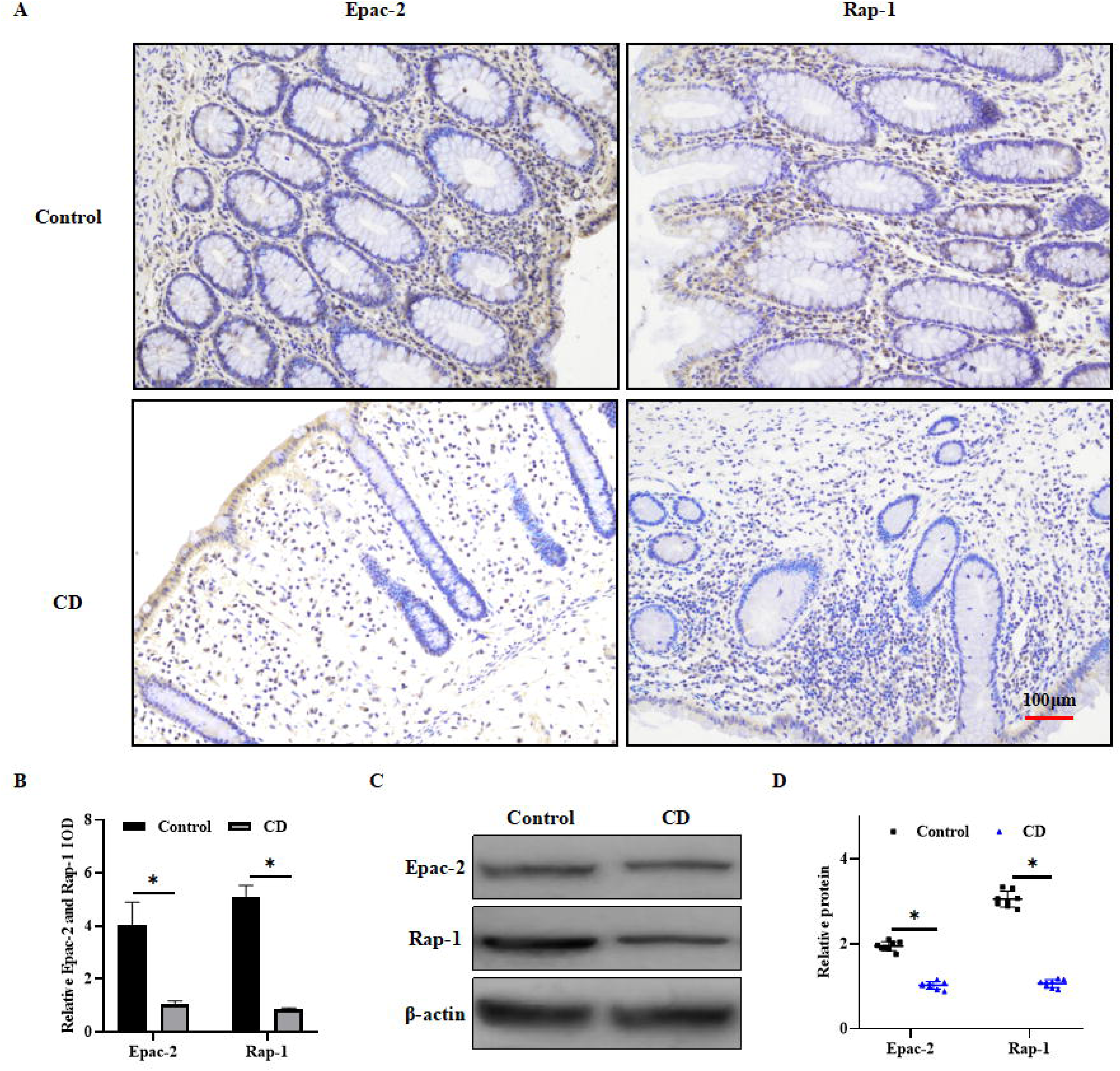
The expressions of Epac-2 and Rap-1 decreased in the colonic tissue of CD patients. (A-B). The IHC analysis reveals Epac-2 and Rap-1 expression were significantly attenuated in colonic tissue of CD patients (n = 12) in contrast to the control patients(n = 16). (C-D) The results of western blotting showed that the levels of Epac-2 and Rap-1 were as well lower in inflammation focus of intestinal tissues from CD patients in contrast to the uninflamed intestines. The data are presented as the relative mean IOD ± SD. * *P* < 0.05.

### 3.5 Rottlerin inhibited LPS-induced FHs 74 int cell apoptosis by activating the Epac-2/Rap-1 pathway

To detect the relationship between the Epac-2/Rap-1 pathway and rottlerin efficacy, we assessed the proliferation and death of LPS-induced FHs 74 int cells by flow cytometry. The proportion of cellular apoptosis in the LPS-induced cells (35.77±4.03%) exceeded that in the control cells (3.07 ± 0.71%) and group treated with LPS and rottlerin (15.29 ± 3.88%). In contrast, treatment with rottlerin and HJC0197 did not significantly reduce LPS-induced apoptosis (34.58 ± 3.79%, **Figure 5A-B**). The results of cell cycle analysis revealed that LPS treatment (74.43±6.32%) led to a significant accumulation of cells in G0/G1 phase in comparison with that in the control group (43.23±3.21%) and the group treated with LPS and rottlerin (59.54 ± 5.65%), but treatment with rottlerin and HJC0197 did not significantly reduce the accumulation of cells in G0/G1 phase (77.38 ± 4.79%, **Figure 5C-D**). The results of Western blotting revealed that the expressions of Epac-2 and Rap-1 were upregulated in the rottlerin team in contrast to the LPS-treated team; nevertheless, the upregulations of Epac-2 and Rap-1 were inhibited by HJC0197 in the group treated with rottlerin (**Figure 5E-F**). Some of these findings proved that the activation of the Epac-2/Rap-1 pathway by rottlerin treatment inhibited LPS-induced FHs 74 int cell apoptosis and maintained proliferation activity.

**Figure 5.**
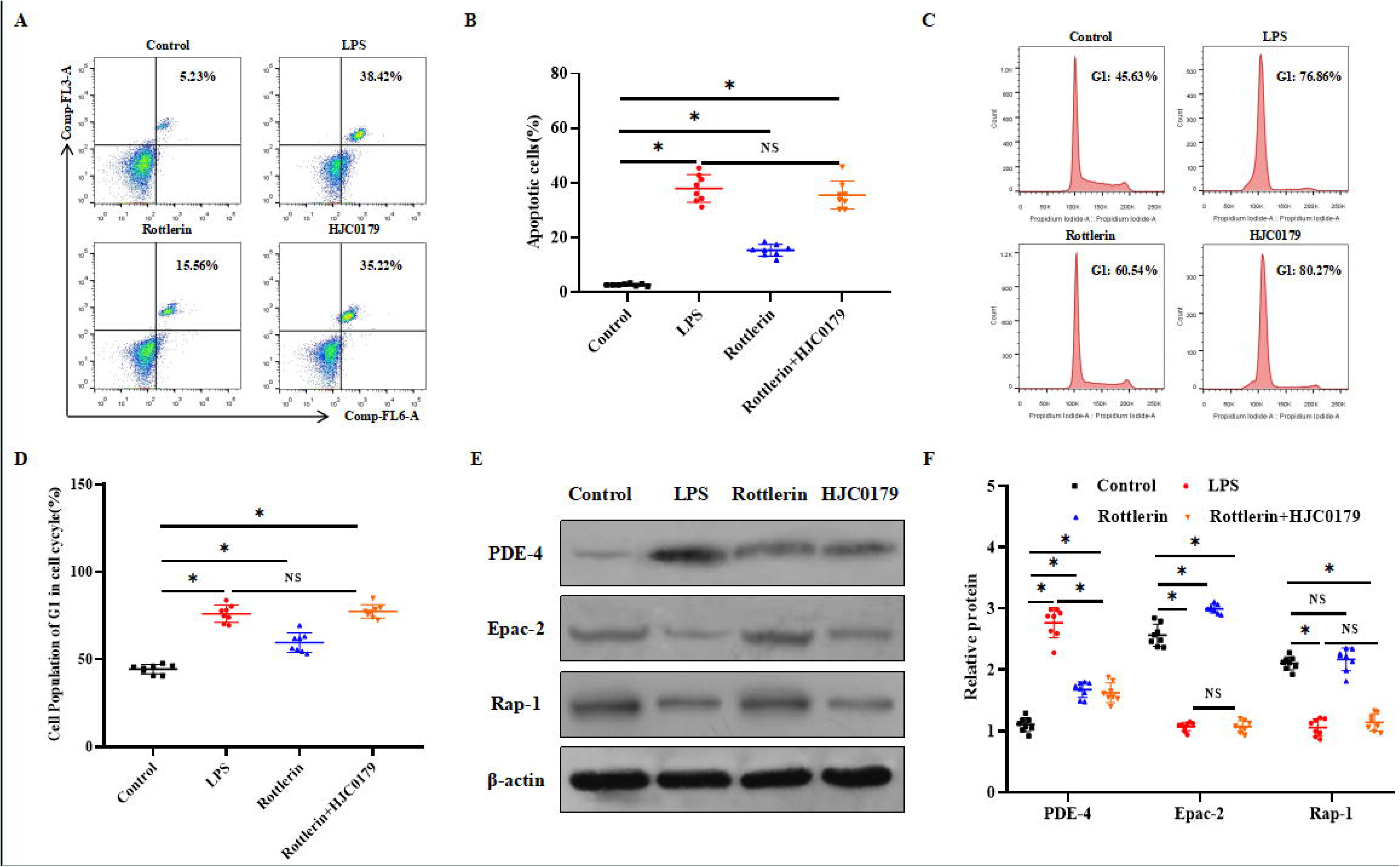
Rottlerin inhibited apoptosis by activating the Epac-2/Rap-1 pathway. (A-B) The proportion of cellular apoptosis in the LPS-treated group was raised in contrast to the control group and rottlerin group, and HJC0197 inhibited the protective effect of rottlerin. (C-D) Cell cycle analysis revealed that rottlerin ameliorated the LPS**-**induced accumulation of cells in G0/G1 phase in contrast to the control group, but that HJC0197 inhibited this trend. (E-F) The datas of Western blot revealed that the levels of Epac-2/Rap-1 were activated in the rottlerin group compared with the LPS-treated group. PDE-4 showed the opposite trend, but HJC0197 inhibited the activation of Epac-2/Rap-1 by rottlerin without changing PDE-4 expression. The detection were executed 3 times separately (8 mice of every group), a representative result is given. The datas are appeared as the means ± SD (\**P* < 0.05).

### 3.6 Rottlerin upregulated the levels of TJ proteins in LPS-induced FHs 74 int cells by activating the Epac-2/Rap-1 pathway

In order to investigatet whether the distributionand and levels of TJ proteins have changed due to the activation of the Epac-2/Rap-1 pathway, immunofluorescence and Western blotting were performed to detect the levels of occludin and ZO-1 protein in the FHs 74 int cells. The levels of ZO-1 and occludin in FHs 74 int cells were obviously upregulated in the rottlerin group in contrast to the LPS-treated group; however, the distribution of occludin and ZO-1 was not recovered in the group treated with LPS, rottlerin and HJC0197 group in contrast to the LPS-treated group. However, no remarkable difference in the protein levels were observed between the rottlerin team and WT mice (**Figure 6A-B**). This trend was confirmed again by the results of western blotting, and the expression of occludin and ZO-1 was upregulated in the rottlerin group in contrast to the LPS-treated group; but the levels were not upregulated in the group treated with LPS, rottlerin and HJC0197 (**Figure 6C-D**). Some of these findings proved that the activation of the Epac-2/Rap-1 pathway by rottlerin treatment upregulated the levels of TJ proteins when the FHs 74 int cells were damaged by LPS.

**Figure 6.**
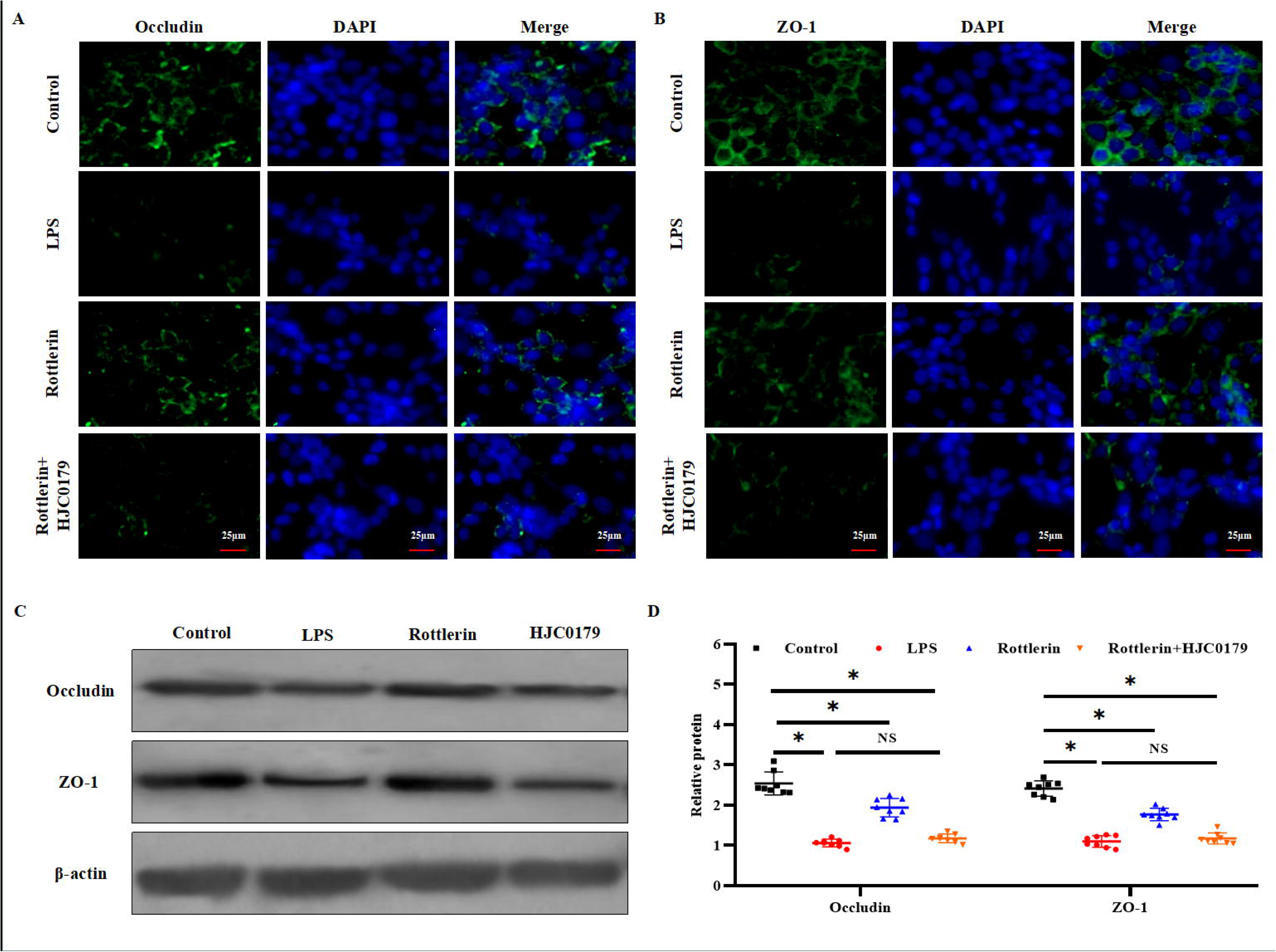
Rottlerin upregulated the levels of TJ proteins by activating the Epac-2/Rap-1 pathway. (A-B) IF revealed that the distributions of occludin and ZO-1 (green) was enhanced in the rottlerin group compared with the LPS**-**treated FHs 74 int cells but that HJC0197 inhibited this trend. The nuclei was stained by DAPI (blue). (C-D) The levels of occludin and ZO-1 in the rottlerin group were clearly raised in contrast to the LPS**-**treated FHs 74 int cells and were similar to those in the control group, but HJC0197 inhibited the effect of rottlerin. The detection were executed 3 times separately (8 mice of every group), a representative result is given. The datas are appeared as the means ± SD (\**P* < 0.05).

## 4. Discussion

For all we know, this research was the first to prove that rottlerin ameliorated DSS-induced colitis in WT mice. The concrete outcomes were as follows: (1) rottlerin improved DSS-induced CD-like colitis via upregulation of the TJ proteins levels; (2) rottlerin preserved the barrier function of DSS-treated mice by reducing the apoptosis rate of epithelial cells; and (3) attenuating PI3K/Akt signaling and activating the Epac-2/Rap-1 pathway in the intestinal mucosa the DSS-induced attenuation of TJ proteins in mice.

This study showed that rottlerin possessed a significant protection function on DSS-induced colitis in WT mice. In the process of intestinal inflammation in patients with CD, proinflammatory factors consisted of TNF-α, IL-1β and IL-17A were highly expressed during intestinal mucosal barrier injury ^[28]^. Our results suggested that the function of rottlerin depended on decreased inflammation scores, DAI values and expression of proinflammatory cytokines. These achievements urge us to further our research, and we discovered that rottlerin treatment reduced the apoptosis rate of epithelial cells in DSS-treated mice. In addition, the activation of Bcl-2 has a powerful antiapoptotic effect, and the ratio of Bcl-2 and Bax is an important additional contributor affecting the balance of apoptosis ^[29, 30]^. We found that treatment with rottlerin increased the level of Bcl-2 in DSS-treated group and decreased the level of Bax. Among many kinds of caspases, caspase-3 plays a key role in promoting the death of cells ^[31]^. Correspond to our observations, the datas showed that the expression of caspase-3 was reduced in the rottlerin group. In order to examine the resist inflammation effects of rottlerin and its mechanisms, we analyzed functional changes in the intestinal barrier related to the pathophysiologic mechanisms of CD. The datas revealed the protective effect of rottlerin in DSS-treated mice was partially achieved by increasing the levels of occludin and ZO-1 in the colonic tissue. Impairments in intestinal epithelial barrier function increased the permeability of intestine and promoted intestinal inflammation in the progression of CD ^[32]^. Our datas indicated that serum glucan binding levels in the rottlerin group and the probability of bacterial transfer to liver and MLN were significantly reduced, further demonstrating the protective effect of rosin on intestinal barrier function. These results suggest that rottlerin has a strong protection function on colitis.

The protection function of rottlerin on the epithelial barrier in DSS-treated group was special encouraging and confirmed the view that rottlerin had latent curative effects on colitis. We found that the administration of rottlerin attenuated PI3K/Akt signaling and activated the Epac-2/Rap-1 pathway. These achievements prompted us to proceed examining potential changes in CD patients ^[33]^. We found that the levels of Epac-2 and Rap-1 in the colonic tissue of CD patients was dramatically reduced in contrast to the control patients. We evaluated the effects of the Epac-2/Rap-1 pathway on the regulation of key TJ proteins and found that rottlerin inhibited apoptosis and promoted TJ protein expression in LPS-induced FHs 74 int cells ^[34]^. In addition, the latest researches have suggested that the signalpathway had a key protective effect on barrier function ^[35, 36]^. Some of these findings proved that the effect of rottlerin on TJ protein expression in DSS-treated mice.

It is undeniable that there are some limitations in our research. For example, our datas indicate rottlerin can ameliorate the inflammatory response and ameliorate colitis via protecting the integrity of the intestinal barrier structure; however, rottlerin can also ameliorate colitis in other ways. The attenuation of PI3K/Akt and the upregulation of the Epac-2/Rap-1 pathway may in part illustrate the protective effect of rottlerin on the function of barrier and its mechanism of action, but other signal channels also should be discussed. As we said before, rottlerin perhaps had various biological activities ^[37]^.

In summary, this research partially reveales that the rottlerin treatment can improve pathological process associated with colitis in DSS-treated group by increasing the levels of TJ proteins and decreasing the apoptosis rate of epithelial cells. Rottlerin may play an active role by reducing PI3K/Akt signaling and activating the Epac-2/Rap-1 pathway in the intestinal mucosa. All the datas reveal rottlerin has a significant protection function on \colitis and offer a novel option for maintenance drug.

## Author contributions

X. Song and L. Zuo: Conceptualization, Methodology, Validation, Resources, Writing - Original Draft, Funding acquisition, Project administration. J. Hu: Conceptualization, Writing - Review & Editing, Funding acquisition, Supervision. L. Wang, Z. Zhu, J. Tao, Y. Jiang, X. Wu, Z. Wang, J. nian and Y. Wang: Data Curation, Formal analysis, Investigation. P. Xiang, X. Zhang, H. Zhao, L. Yu and J. Li : Methodology, Resources, Funding acquisition, Visualization. All authors read and approved the final manuscript.

## Acknowledgments

This work was supported partly by funding from the First Affiliated Hospital of Bengbu Medical College Science Fund for Outstanding Young Scholars (2019byyfyyq02, 2019byyfyyq08), the First Affiliated Hospital of Bengbu Medical College Science Fund for Distinguished Young Scholars (2019byyfyjq01), the Natural Science Foundation of Anhui Province (1808085QH237, KJ2018A1001, KJ2019A0328), the National Natural Science Foundation of China (81700476 and 81500421), the Natural Science Foundation of Bengbu Medical College (BYKY1721ZD, BYKY1888) and the Technology Development Foundation of First Affiliated Hospital of Bengbu Medical College (Byyfykj201802).

## References

[1] Lim WC, Wang Y, MacDonald JK, Hanauer S. Aminosalicylates for induction of remission or response in Crohn’s disease. Cochrane Database Syst Rev 2016;7:CD008870.

[2] Iheozor-Ejiofor Z, Gordon M, Clegg A, Freeman SC, Gjuladin-Hellon T, MacDonald JK, et al. Interventions for maintenance of surgically induced remission in Crohn’s disease: a network meta-analysis. Cochrane Database Syst Rev 2019;9:CD013210.

[3] Maoz A, Dennis M, Greenson JK. The Crohn’s-Like Lymphoid Reaction to Colorectal Cancer-Tertiary Lymphoid Structures With Immunologic and Potentially Therapeutic Relevance in Colorectal Cancer. Front Immunol 2019;10:1884.

[4] de Bruyn M, Vermeire S. NOD2 and bacterial recognition as therapeutic targets for Crohn’s disease. Expert Opin Ther Targets 2017;21:1123–1139.

[5] Pêgo B, Martinusso CA, Bernardazzi C, Ribeiro BE, de Araujo Cunha AF, de Souza Mesquita J, et al. Schistosoma mansoni Coinfection Attenuates Murine Toxoplasma gondii-Induced Crohn’s-Like Ileitis by Preserving the Epithelial Barrier and Downregulating the Inflammatory Response. Front Immunol 2019;10:442.

[6] Song X, Li J, Wang Y, Zhou C, Zhang Z, Shen M, et al. Clematichinenoside AR ameliorated spontaneous colitis in Il-10-/-mice associated with improving the intestinal barrier function and abnormal immune responses. Life Sci 2019;239:117021.

[7] Lo BC, Shin SB, Canals HD, Refaeli I, Yu HB, Goebeler V, et al. IL-22 Preserves Gut Epithelial Integrity and Promotes Disease Remission during Chronic Salmonella Infection. J Immunol 2019;202:956–965.

[8] Ramos GP, Papadakis KA. Mechanisms of Disease: Inflammatory Bowel Diseases. Mayo Clin Proc 2019;94:155–165.

[9] Keita ÅV, Lindqvist CM, Öst Å, Cdl M, Schoultz I, Halfvarson J. Gut Barrier Dysfunction-A Primary Defect in Twins with Crohn’s Disease Predominantly Caused by Genetic Predisposition. J Crohns Colitis 2018;12:1200–1209.

[10] Xu CL, Guo Y, Qiao L, Ma L, Cheng YY. Recombinant expressed vasoactive intestinal peptide analogue ameliorates TNBS-induced colitis in rats. World J Gastroenterol 2018;24:706–715.

[11] Assa A, Matar M, Turner D, Broide E, Weiss B, Ledder O, et al. Proactive Monitoring of Adalimumab Trough Concentration Associated With Increased Clinical Remission in Children With Crohn’s Disease Compared With Reactive Monitoring. Gastroenterology 2019;157:985-996.e2.

[12] Viladomiu M, Kivolowitz C, Abdulhamid A, Dogan B, Victorio D, Castellanos JG, et al. IgA-coated E. coli enriched in Crohn’s disease spondyloarthritis promote TH17-dependent inflammation. Sci Transl Med 2017;9.

[13] Dar MI, Mahajan P, Jan S, Jain SK, Tiwari H, Sandey J, et al. Rottlerin is a pan phosphodiesterase inhibitor and can induce neurodifferentiation in IMR-32 human neuroblastoma cells. Eur J Pharmacol 2019;857:172448.

[14] Albrecht W, Unger A, Bauer SM, Laufer SA. Discovery of N-{4-[5-(4-Fluorophenyl)-3-methyl-2-methylsulfanyl-3H-imidazol-4-yl]-pyridin-2-yl}-acetamide (CBS-3595), a Dual p38α MAPK/PDE-4 Inhibitor with Activity against TNFα-Related Diseases. J Med Chem 2017;60:5290–5305.

[15] Suzuki T, Hara H. Quercetin enhances intestinal barrier function through the assembly of zonula [corrected] occludens-2, occludin, and claudin-1 and the expression of claudin-4 in Caco-2 cells. J Nutr 2009;139:965–974.

[16] Li H, Fan C, Feng C, Wu Y, Lu H, He P, et al. Inhibition of phosphodiesterase-4 attenuates murine ulcerative colitis through interference with mucosal immunity. Br J Pharmacol 2019;176:2209–2226.

[17] Birukova AA, Zebda N, Fu P, Poroyko V, Cokic I, Birukov KG. Association between adherens junctions and tight junctions via Rap1 promotes barrier protective effects of oxidized phospholipids. J Cell Physiol 2011;226:2052–2062.

[18] de Rooij J, Zwartkruis FJ, Verheijen MH, Cool RH, Nijman SM, Wittinghofer A, et al. Epac is a Rap1 guanine-nucleotide-exchange factor directly activated by cyclic AMP. Nature 1998;396:474–477.

[19] Wirtz S, Popp V, Kindermann M, Gerlach K, Weigmann B, Fichtner-Feigl S, et al. Chemically induced mouse models of acute and chronic intestinal inflammation. Nat Protoc 2017;12:1295–1309.

[20] Juneja M, Kobelt D, Walther W, Voss C, Smith J, Specker E, et al. Statin and rottlerin small-molecule inhibitors restrict colon cancer progression and metastasis via MACC1. PLoS Biol 2017;15:e2000784.

[21] Zhang H, Peng A, Yu Y, Guo S, Wang M, Wang H. l-Arginine Protects Ovine Intestinal Epithelial Cells from Lipopolysaccharide-Induced Apoptosis through Alleviating Oxidative Stress. J Agric Food Chem 2019;67:1683–1690.

[22] Ramos-Alvarez I, Lee L, Jensen RT. Cyclic AMP-dependent protein kinase A and EPAC mediate VIP and secretin stimulation of PAK4 and activation of Na+,K+-ATPase in pancreatic acinar cells. Am J Physiol Gastrointest Liver Physiol 2019;316:G263–263G277.

[23] Spencer DM, Veldman GM, Banerjee S, Willis J, Levine AD. Distinct inflammatory mechanisms mediate early versus late colitis in mice. Gastroenterology 2002;122:94–105.

[24] Schultz M, Tonkonogy SL, Sellon RK, Veltkamp C, Godfrey VL, Kwon J, et al. IL-2-deficient mice raised under germfree conditions develop delayed mild focal intestinal inflammation. Am J Physiol 1999;276:G1461–1472.

[25] Dong J, Wang H, Zhao J, Sun J, Zhang T, Zuo L, et al. SEW2871 protects from experimental colitis through reduced epithelial cell apoptosis and improved barrier function in interleukin-10 gene-deficient mice. Immunol Res 2015;61:303–311.

[26] Li J, Zuo L, Tian Y, He Y, Zhang Z, Guo P, et al. Spontaneous colitis in IL-10-deficient mice was ameliorated via inhibiting glutaminase1. J Cell Mol Med 2019;23:5632–5641.

[27] Zuo L, Li J, Ge S, Ge Y, Shen M, Wang Y, et al. Bryostatin-1 ameliorated experimental colitis in Il-10-/-Mice by protecting the intestinal barrier and limiting immune dysfunction. J Cell Mol Med 2019;23:5588–5599.

[28] Ma C, Lin W, Liu Z, Tang W, Gautam R, Li H, et al. NDR1 protein kinase promotes IL-17- and TNF-α-mediated inflammation by competitively binding TRAF3. EMBO Rep 2017;18:586–602.

[29] Aghdaei HA, Kadijani AA, Sorrentino D, Mirzaei A, Shahrokh S, Balaii H, et al. An increased Bax/Bcl-2 ratio in circulating inflammatory cells predicts primary response to infliximab in inflammatory bowel disease patients. United European Gastroenterol J 2018;6:1074–1081.

[30] Hu Y, Yagüe E, Zhao J, Wang L, Bai J, Yang Q, et al. Sabutoclax, pan-active BCL-2 protein family antagonist, overcomes drug resistance and eliminates cancer stem cells in breast cancer. Cancer Lett 2018;423:47–59.

[31] Kavanagh E, Rodhe J, Burguillos MA, Venero JL, Joseph B. Regulation of caspase-3 processing by cIAP2 controls the switch between pro-inflammatory activation and cell death in microglia. Cell Death Dis 2014;5:e1565.

[32] Westbrook AM, Szakmary A, Schiestl RH. Mouse models of intestinal inflammation and cancer. Arch Toxicol 2016;90:2109–2130.

[33] Oshima T, Miwa H. Gastrointestinal mucosal barrier function and diseases. J Gastroenterol 2016;51:768–778.

[34] Angelow S, Zeni P, Höhn B, Galla HJ. Phorbol ester induced short- and long-term permeabilization of the blood-CSF barrier in vitro. Brain Res 2005;1063:168–179.

[35] Spindler V, Peter D, Harms GS, Asan E, Waschke J. Ultrastructural analysis reveals cAMP-dependent enhancement of microvascular endothelial barrier functions via Rac1-mediated reorganization of intercellular junctions. Am J Pathol 2011;178:2424–2436.

[36] Baumer Y, Spindler V, Werthmann RC, Bünemann M, Waschke J. Role of Rac 1 and cAMP in endothelial barrier stabilization and thrombin-induced barrier breakdown. J Cell Physiol 2009;220:716–726.

[37] Kim YH, Kim D, Hong AR, Kim JH, Yoo H, Kim J, et al. Therapeutic Potential of Rottlerin for Skin Hyperpigmentary Disorders by Inhibiting the Transcriptional Activity of CREB-Regulated Transcription Coactivators. J Invest Dermatol 2019;139:2359-2367.e2.

